# From citizen to scientist: evaluating ant observation accuracy on a Mediterranean archipelago

**DOI:** 10.1101/2025.02.08.637263

**Authors:** Javier Arcos

## Abstract

Citizen science platforms, such as iNaturalist, have become invaluable tools for biodiversity monitoring, allowing non-experts to contribute photo-based observations that are then identified by the user community. However, one of the major challenges of these platforms concerns the accuracy of these identifications, especially for taxa requiring specialized knowledge or more sophisticated means of identification. In this study, I assess the reliability of ant identifications in the Balearic Islands by comparing community-generated identifications with expert validations across multiple taxonomic levels. Based on 300 iNaturalist observations, I analyze the influence of user experience, AI-generated suggestions, image quality, and community participation on identification accuracy. The results indicate that species-level accuracy was 69.91%, increasing to 90.91% at the genus level and 99% at the family level. Experienced users significantly improved accuracy, while early-stage identifications at finer taxonomic resolutions increased the likelihood of correct consensus. AI-assisted identifications performed well for frequently recorded species but struggled with underrepresented taxa. Surprisingly, photo quality had minimal impact, as common species were often identifiable even from low-resolution images. Additionally, the dataset documented exotic species, including the first record of the *Formica rufibarbis* complex in Mallorca. These findings highlight both the strengths and limitations of citizen science in taxonomic research and emphasize the need for strategies to enhance data reliability for conservation and biodiversity monitoring.

## Introduction

Citizen science has increasingly become a powerful tool in ecological and environmental research, enabling the collection of vast amounts of data that would be challenging, if not impossible, to gather through traditional methods (Kosmala et al., 2016). One notable example is iNaturalist, an online platform that allows citizens, enthusiasts, and professionals to participate in monitoring biodiversity by submitting observations on different taxa. These observations, often accompanied by photographs or audio recordings, are then collaboratively identified by a diverse community of users (Campbell et al., 2023). This participatory approach not only democratizes science but also facilitates global-scale research, fostering collaboration between non-specialists and experts (Bonney et al., 2014; Fraisl et al., 2022). Over time, iNaturalist has proven invaluable in documenting species distributions, tracking ecological changes, and improving the early detection of invasive species (Di Cecco et al., 2021; Rosa et al., 2022; Encarnação et al., 2021; Mesaglio & Callaghan, 2021). Despite its many strengths, the reliability of the data collected through iNaturalist remains a critical challenge, particularly in terms of its suitability for use in scientific publications and research. Ensuring the accuracy, consistency, and usability of these citizen science contributions is essential to maximize their value in ecological studies and environmental monitoring (Kosmala et al., 2016; Koo et al., 2022).

The Balearic Islands, a Mediterranean archipelago that hosts a well-documented ant fauna, supported by extensive fieldwork conducted in previous decades, with 67 known species (Arcos & García, 2024), provides an excellent case study for evaluating the reliability of ant identifications on iNaturalist. This region features a diverse ant community, including both native and exotic species, making it a suitable setting to test the accuracy of citizen science data. Since ants are one of the most common insect groups in the world, easily observed and usually exhibit subtle morphological differences that can frequently lead to misidentifications, their study should therefore be ideal for looking into the accuracy of citizen science data.

Overall, this study aims to assess the validity of ant identifications on iNaturalist within the Balearic Islands by analyzing a medium-sized sample of unstructured, opportunistic observations. It compares community-contributed identifications with expert-verified ones to evaluate how reliably citizen scientists can identify ant species in this region. The study also investigates the influence of factors such as user experience, the quality of photographs submitted, and the effectiveness of AI tools in the identification process. By addressing these challenges, the findings aim to further improve the reliability of citizen science platforms like iNaturalist, providing useful data to monitor and protect biodiversity.

## Materials and Methods

### Data collection

Observations of ants were retrieved from iNaturalist using the platform’s API. The form was set to include only records from the family Formicidae within the Balearic Islands, which comprise Mallorca, Menorca, Eivissa, Formentera, and some smaller islets. The dataset was downloaded on October 5, 2024, and included variables provided by the platform such as observation ID, location, date, observer name, quality grade, and suggested taxon. A total of 326 observations were initially retrieved. To enhance the dataset, additional variables not included in the standard iNaturalist export’s form were retrieved in August 2024 using a custom script developed by the first author using Python. These variables included the names and counts of identifiers involved in each observation, the total number of identifications, the presence or absence of AI-generated suggestions and their proposed taxa, or the taxonomic category agreed upon by the community. These variables were compiled into a single database for analysis.

### Definitions and data preparation

Observations on iNaturalist are classified into three quality grades—Casual, Needs ID, and Research Grade—based on criteria such as data completeness, photo quality, and user consensus (Campbell et al. 2023). In this study, the Community Taxon, which represents the taxonomic level with at least two-thirds agreement among contributors, was used as the primary identifier for analyses. For the analysis, precision was defined as the consensus or disagreement at the taxonomic category proposed by the expert for the community ID and the ID of the taxonomic resolution applied to the Expert ID. Conversely, accuracy was defined considering Community ID and Expert ID at various taxonomic levels: species, genus, subfamily and family and assessing concordance at successively higher hierarchical levels. Comparisons were possible whenever one of the paired IDs contained a value for the level of interest. For example, if an observation did not have a value at the species level for the Community ID because the ID applied to a higher taxonomic rank but the Expert ID did contain a value, this was scored as a mismatch.

The dataset was carefully filtered to ensure validity and relevance. Observations graded as Casual were excluded if they lacked visible ants, had misplaced geographical coordinates, or were of captive specimens. Observations labeled as Needs ID were excluded if they lacked diagnostic traits to be treated as ants. Records with insufficient identifications to generate a Community Taxon were also removed. A total of 26 observations were excluded, including five Casual observations, two Needs ID records, and 19 observations with insufficient IDs, leaving 300 valid observations for analysis.

### Expert validation

Each observation was reviewed by the author, with experience in both Mediterranean ant taxonomy and citizen science validation. The expert’s identification (*taxonomic_unit*) was considered the ground truth for evaluating the precision and accuracy of community-generated. To ensure unbiased identifications, the expert reviewed the observations blindly, accessing only the photo links without contextual information such as location or previously suggested IDs. The Community Taxon was then compared with the Expert ID across multiple taxonomic levels, including family, subfamily, genus, and species, inferred upwards in the taxonomic ranking for each taxonomic unit.

### Variables for analysis

To examine factors influencing identification accuracy, the study included variables related to observation quality, community dynamics, and AI suggestions (Table 1). The total number of identifications submitted for each observation, the number of unique contributors, and the counts of agreements and disagreements among identifiers were analyzed. AI-related variables included the presence or absence of an AI-generated suggestion, the taxon proposed by the AI, and whether the species was included in the AI training dataset. Photo quality was categorized as low, medium, or high based on image resolution and the visibility of diagnostic traits. Identification difficulty, scored from one to three, reflected the inherent complexity of distinguishing the species.

**Table 1.**
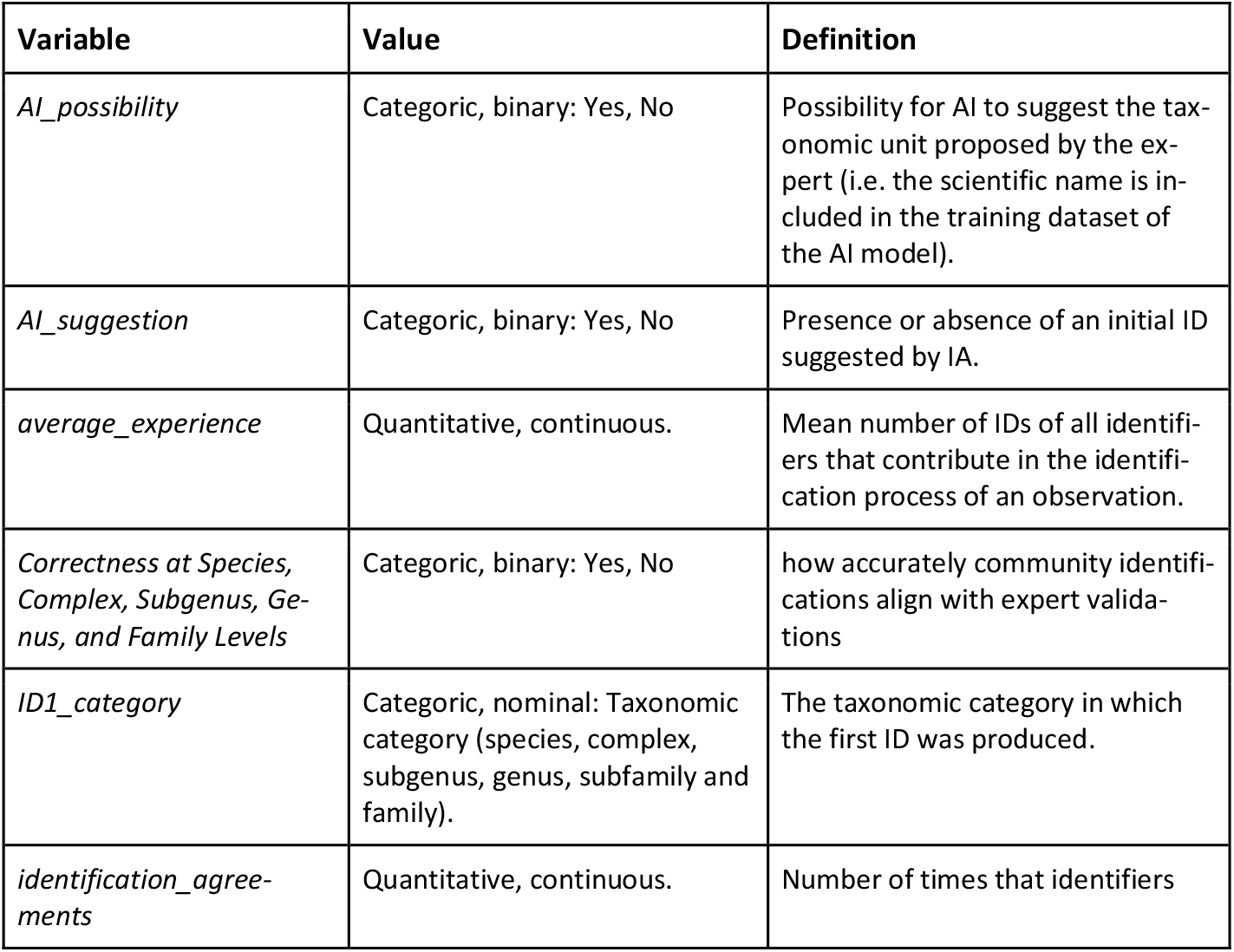
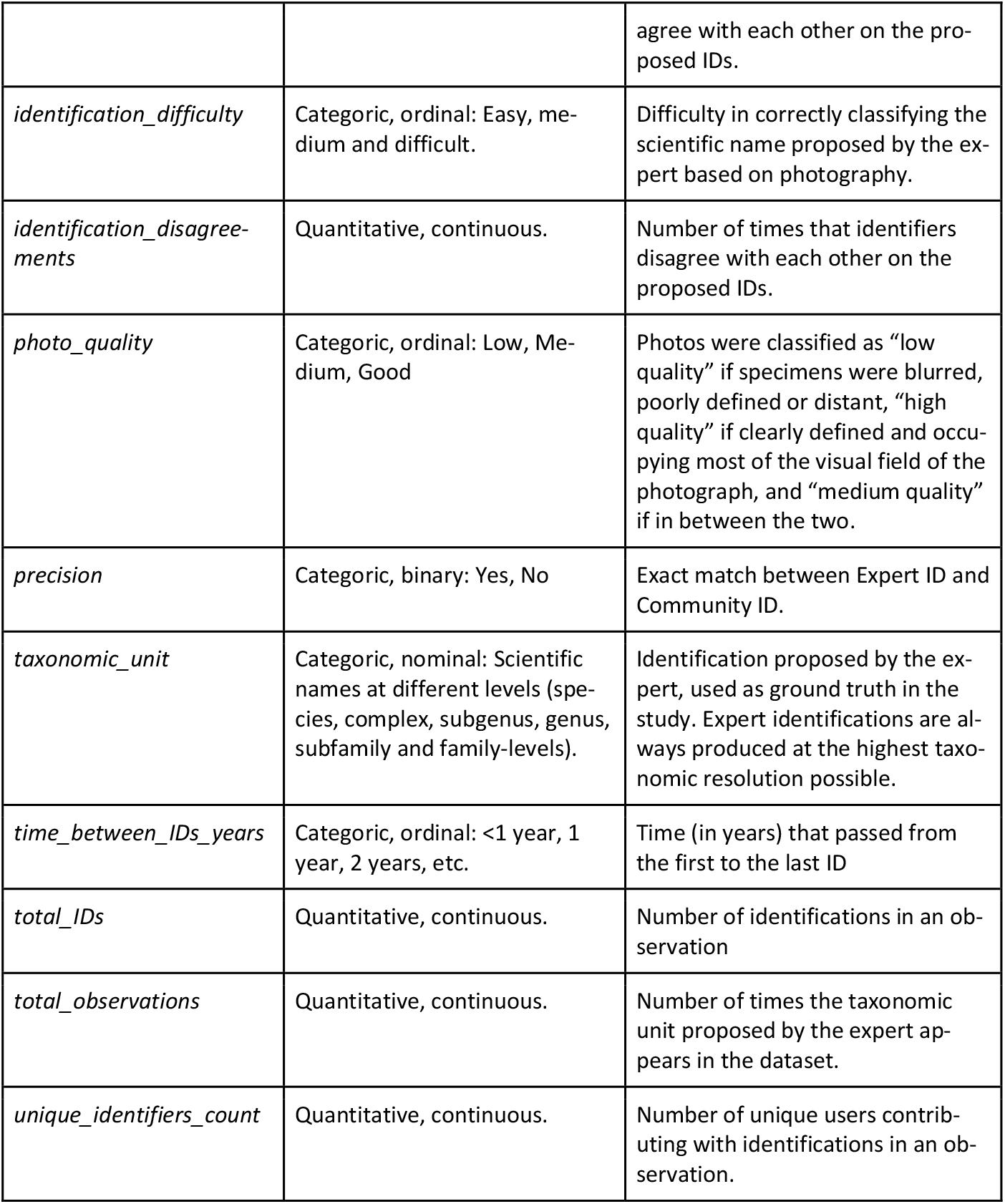
Variables used in the study, with their values and definition.

### Statistical and analytical methods

The primary dependent variable was precision and accuracy, which evaluated whether the Community Taxon matched the Expert Decision (*taxonomic_unit)*. Identification accuracy was calculated as the proportion of correct identifications at each taxonomic level, including family, genus, and species. Statistical analyses were conducted to evaluate the influence of independent variables on identification accuracy. A correlation matrix was used to study associations between variables. Machine learning approaches, including a Random Forest classifier, were used to predict identification accuracy and to identify the relative importance of each variable. To ensure robust model validation, the dataset was divided into training and test subsets using a 70:30 split. Some ordinal, categorical variables such as *photo_quality* were transformed using One-Hot Encoding to avoid artificial lineality when testing correlations with other variables. Chi-square tests were used to compare categorical variables, such as the accuracy of identifications at different taxonomic levels, while paired t-tests assessed differences between experienced users and community averages. Logistic regression models were applied to evaluate the effects of variables such as photo quality, identification difficulty, and the number of contributors on the likelihood of a correct identification.

## Results

### 1. Overview of the iNaturalist Ant Observations Dataset

The iNaturalist Ant Observations Dataset includes 300 valid observations of ants from the Balearic Islands, contributed by 113 unique observers. The records are categorized as follows: 168 are “Research Grade,” 132 are “Needs ID” and 5 are “Casual”. Overall, a total of 1,353 IDs were contributed by 193 different unique identifiers, with 18.54% of these identifiers responsible for about 80% of the total number of IDs, emphasizing a strong contribution of a few highly active users. In fact, only the top 7 users are needed to reach nearly half (49.52%) of all IDs, with the top two identifiers producing as much as 178 and 107 IDs, correspondingly. The average number of IDs per observation was 4.51 ± 1.67. Most of the observations included between 3 and 5 IDs (Graph 1). The greatest number of IDs for any observation was 10 (n=1). The top 15 observers produced 50.0% of all 300 observations. For 65.33% of the observations (n=196), AI-generated initial ID suggestions were available, showing its successful implementation on the platform. The observations in the dataset ranged from 2017 to 2024 (Graph 2), but 87.08% of them took place from 2021 forward, demonstrating the increasing popularity of citizen science platforms in recent years. Some of the ants belonged to photographs as old as 2011 and 2012, but they were uploaded starting from 2017. For 59.33% of the observations, less than one year has passed from the first ID to the last.

The dataset consists of 26 expert-validated taxa, including 18 species and 8 taxonomic units representing at least one additional species not identifiable at a finer level via photography: complexes *Aphaenogaster sardoa, Formica rufibarbis, Monomorium salomonis* and *Tetramorium caespitum*, genus *Plagiolepis, Solenopsis* and *Tapinoma*, and subgenus *Lasius*. They are grouped in 17 genera, 3 sub-families, and 1 family (Formicidae) (Graph 3). On average, 2.25 species were newly identified each year. The top 10 species of the iNaturalist Ant Observations Dataset accounted for about 95.67% of all the observations. These were mostly easily recognizable and abundant ants, including *Messor bouvieri* at 22.67%, *Crematogaster scutellaris* at 18.67%, *Pheidole pallidula* at 7.00%, *Linepithema humile* at 6.00%, and *Messor ibericus* at 4.33%. At the genus level, *Messo*r had the highest number of observations with 32.66%, followed by *Crematogaster* with 22.90% and *Camponotus* with 7.74%. Subfamily Myrmicinae gathered 74.83% of the observations, followed by Formicinae (16.78%) and Dolichoderinae (8.39%). The photo quality distribution of the dataset showed that images were mostly of low (49.00%) and medium quality (47.00%), with only 4.00% categorized as “high quality” (Image 1). The difficulty to identify the taxonomic unit proposed by the expert was rated as “medium” at 55.33%, followed by “easy” (38.33%) and “difficult” (6.33%).

**Image 1.**
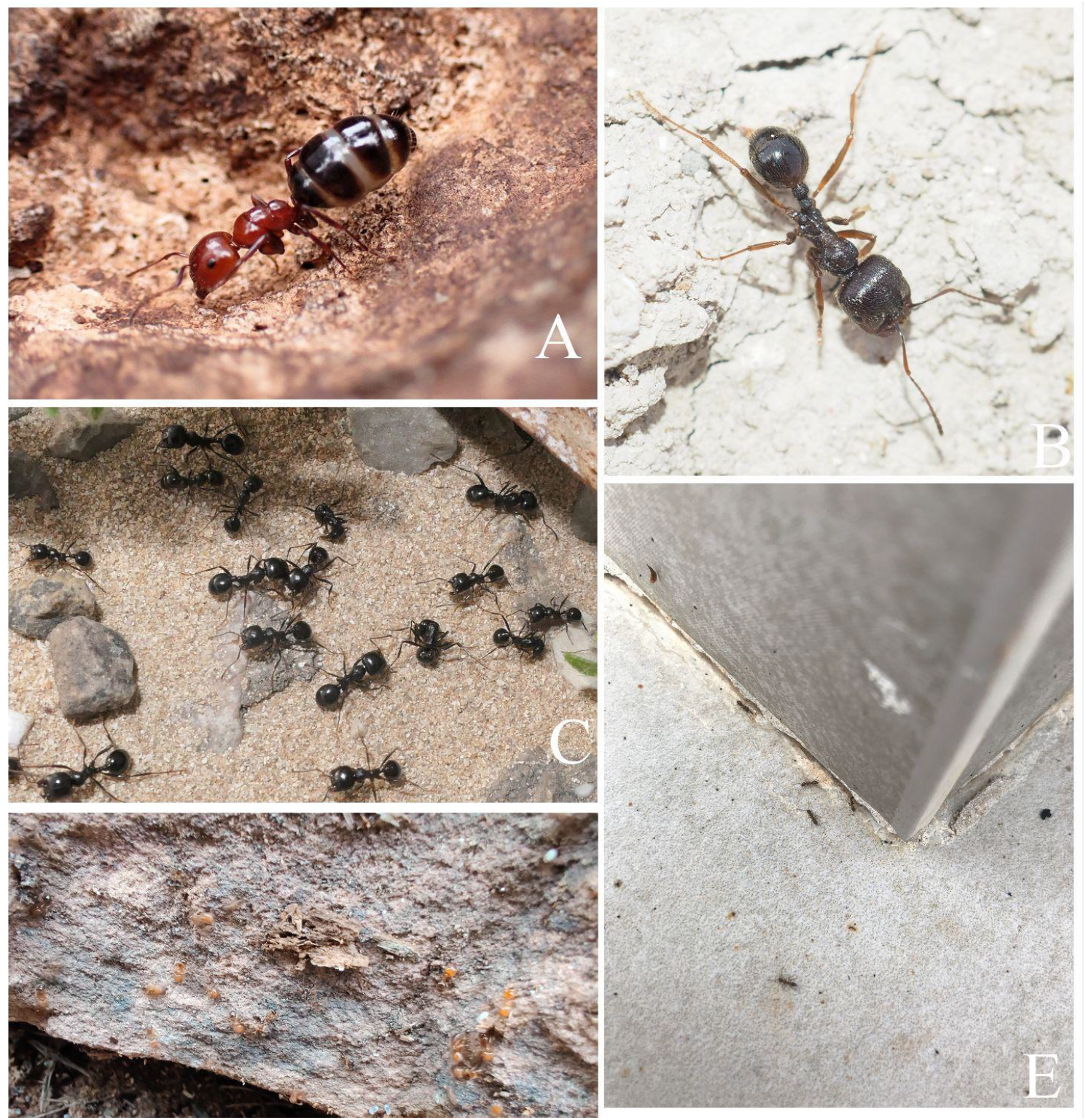
Common species in the dataset and examples of variation in photo quality. A) Good quality photo of a worker of *Camponotus ruber* (© velinr). B) Good quality photo of a *M. ibericus* worker (© fauna_mirifica). C) Medium quality photo of *M. bouvier*i worker (© Jon J. Laysell). D) Low quality photo of soldiers and minors of *P. pallidula* (© geodroid). E) Low quality photo of workers of *L. humile* (© ocean_explorers).

**Graph 1.**
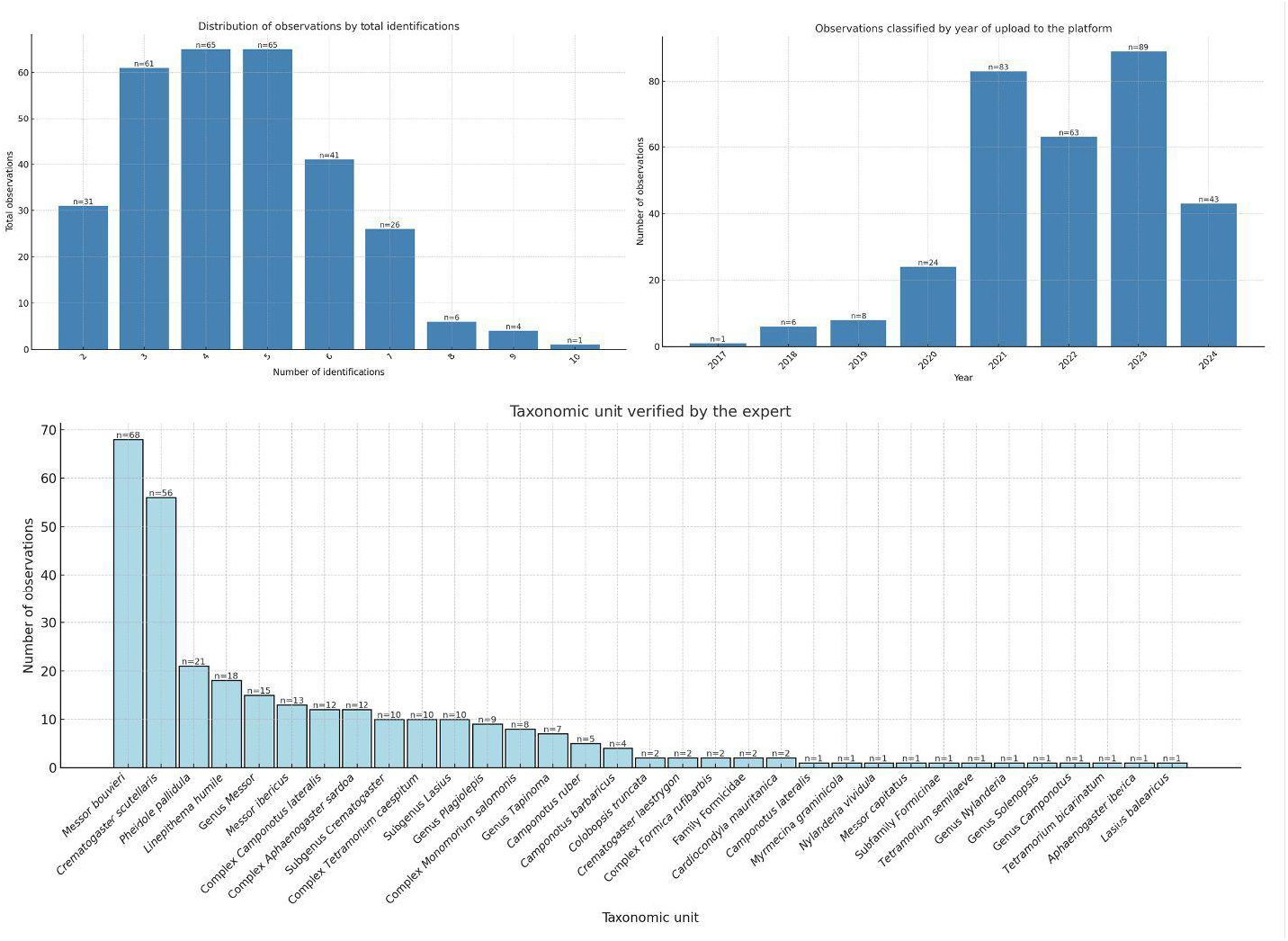
(top left). Distribution of observations according to their total number of IDs. Graph 2 (top right). Observations classified by the year they were uploaded to the platform. Graph 3 (bottom). Frequency of observations for each taxonomic unit in the dataset.

### 2. How important is each analyzed variable?

Thirteen variables were analyzed to assess their contribution to the precision and accuracy of ant observations on iNaturalist (Table 1). To analyze the associations among these variables, a correlation matrix was generated by merging numerical data with encoded categorical variables (Graph 4). The analysis exposed several significant associations (Table 3). Notably, the strongest positive correlations were observed between *total_IDs* and *unique_identifiers_count* (r = 0.93), between *total_IDs* and *num_identification_agreements* (r = 0.65), and between *unique_identifiers_count* and *num_identification_agreements* (r = 0.75). This pattern was expected because a higher number of identifications per observation generally implies greater participation by identifiers, which in turn leads to more agreement. The total number of observations (*total_observations*) and the possibility of AI-suggestion (*AI_possibility*) also demonstrated a good correlation (r= 0.47); this is mostly explained because the AI-recognition model of iNaturalist is often trained with the most frequent species uploaded to the platform.

Negative correlations were observed between *time_between_IDs_years* and *unique_identifiers_count* (r = -0.49), between *time_between_IDs_years* and *total_IDs* (r = -0.49), between *time_between_IDs_years* and *num_identification_agreement*s (r = -0.36), and between *time_between_IDs_years* and *average_experience* (r = -0.43). These negative associations suggest that older observations tend to have fewer identifiers and identifications, resulting in lower counts of agreements and accumulated user experience. This trend likely reflects the lower popularity of the platform in earlier years, when participation was limited and observations were probably never revisited by newer users. Consequently, these older contributions may also have been made by less experienced users who did not continue active participation in subsequent years.

Subsequently, a Random Forest analysis —renowned for its robustness in handling correlated variables— was performed independently to evaluate the relevance of each variable. The resulting metrics are presented in Tables 4 and 5. The Random Forest model achieved an accuracy of 73.33% for predicting precision of community consensus, demonstrating good performance in estimating precision within the dataset. For predictions of accuracy at the species, subgenus, genus, and family levels, the model performed better, with accuracy rates ranging from 84.09% to 100%. These results highlight the value of certain variables across the different models of precision and accuracy. Overall, the most influential variable in the models was *average_experience*, underscoring the key role of experienced users in the identification process. The second most relevant variable was *total_observations*, which represents the number of observations per taxon in the dataset or the resolution of the initial identification. Other notable variables included *ID1_category*, reflecting the category assigned in the initial identification. In addition, *total_IDs, identification_agreements*, and *unique_identifiers_count* demonstrated significant predictive value, indicating that the number of identifications per observation—as well as the number of identifiers and their agreements—are important factors. Beyond these top six variables, additional contributors such as *time_between_ID_years*, AI-related variables and *photo_quality* played minor roles.

**Graph 4.**
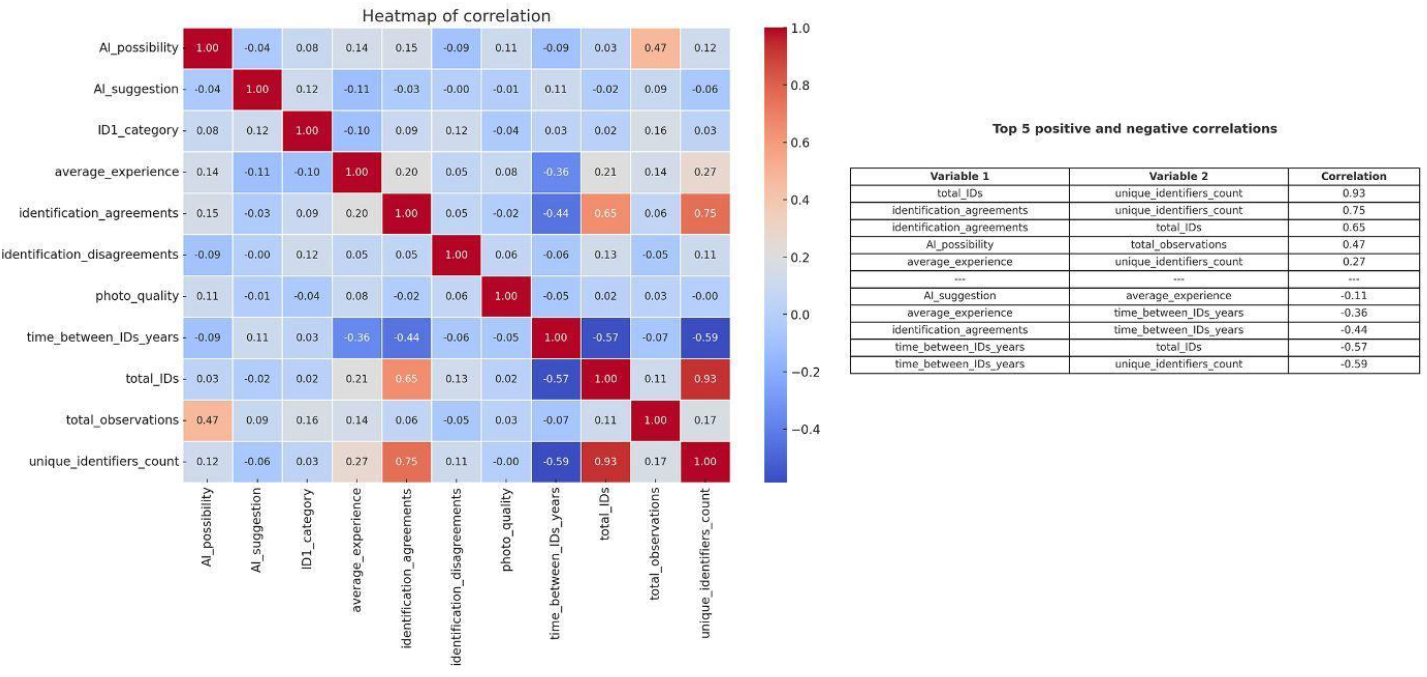
(left). Correlation matrix of analyzed variables. Table 3 (right). Numerical values of the top 5 most positive and negative correlations found.

**Table 4.**
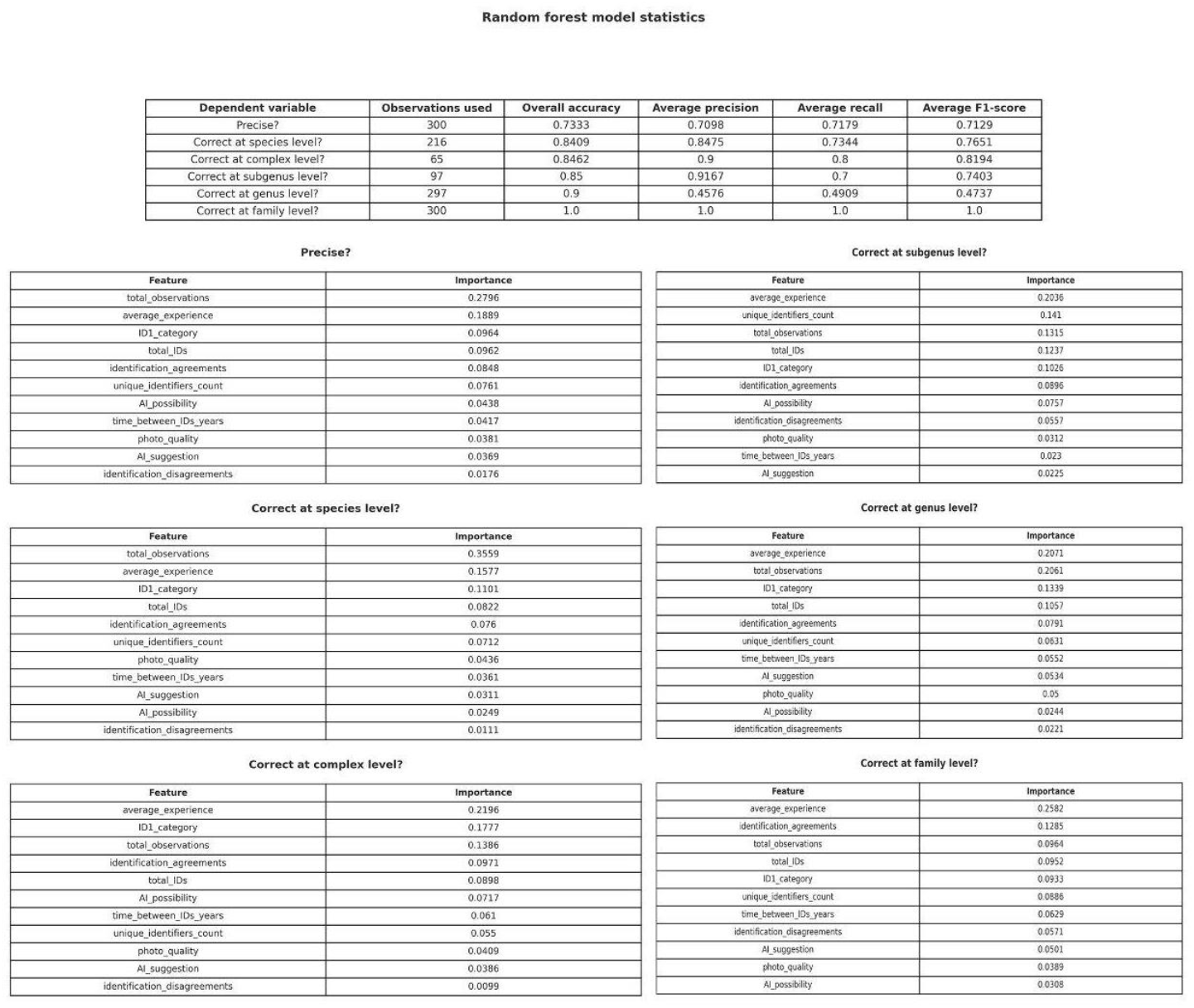
Random Forests statistics and importance of variables for each model.

**Table 5.** Average importance of each variable across all Random Forest models, with standard deviation, as well as maximum and minimum values.

| Feature | Average importance | Standard deviation | Max - Min |
| --- | --- | --- | --- |
| average_experience | 0.2058 | 0.0304 | 0.2582 - 0.1577 |
| total_observations | 0.2014 | 0.0912 | 0.3559 - 0.0964 |
| ID1_category | 0.1190 | 0.0294 | 0.1777 - 0.0933 |
| total_IDs | 0.0988 | 0.0132 | 0.1237 - 0.0822 |
| identification_agreements | 0.0925 | 0.0175 | 0.1285 - 0.0760 |
| unique_identifiers_count | 0.0825 | 0.0282 | 0.1410 - 0.0550 |
| time_between_IDs_years | 0.0466 | 0.0144 | 0.0629 - 0.0230 |
| AI_possibility | 0.0452 | 0.0212 | 0.0757 - 0.0244 |
| photo_quality | 0.0404 | 0.0057 | 0.0500 - 0.0312 |
| AI_suggestion | 0.0388 | 0.0106 | 0.0534 - 0.0225 |
| identification_disagreements | 0.0289 | 0.0199 | 0.0571 - 0.0099 |

### 3. How reliable are citizen scientists in identifying ants on iNaturalist?

The accuracy for ant identifications on iNaturalist was 69.91% at the species level (216 valid comparisons), 90.91% at the genus level (297 comparisons), 96.31% at the subfamily level (298 comparisons), and 99% at the family level (300 comparisons). The overall precision, however, plunged to 64.0% for all 300 observations. This decrease is mainly explained due to the higher taxonomic resolution of the expert compared to the community’s IDs, as illustrated in the heatmap in Graph 5. For example, many species-level IDs made by the expert did not match the genus-level IDs provided by the community. Expert IDs were mainly at the species level (66.33%), higher than the community’s (57.00%). The expert also made more species-complex-level IDs (14.67%) than the community (7.67%). The community, on the other hand, had a higher proportion of genus-level IDs (25.33%) compared to the expert’s (11.33%) and made more identifications at the subfamily (5.00%) and family levels (2.33%) than the expert, who identified at these levels far less frequently (0.33% for subfamily and 0.67% for family). Precision varied between 69% and 100% in moderately to commonly recorded species, while uncommonly recorded species had between 0% and 100% agreement. In conclusion, the analysis indicates that accuracy decreases as taxonomic resolution becomes finer, which helps explain why overall precision is only moderate given the expert’s tendency to identify at a higher resolution.

**Graph 5.**
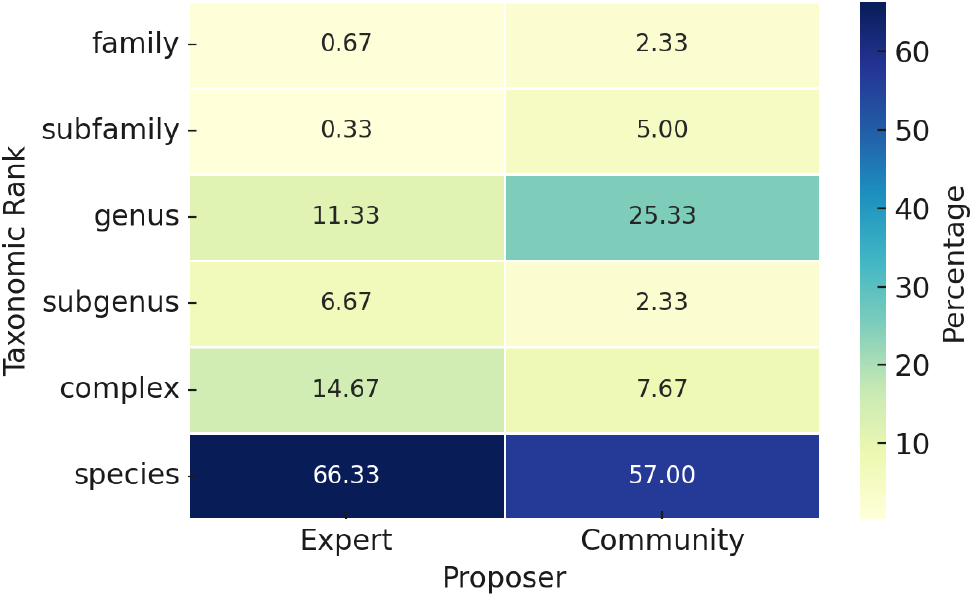
Heatmap showing the percentage comparison of categories proposed by the expert versus the community. The values are displayed as percentages, with different shades indicating the distribution across taxonomic levels.

### 4. Does experience matter? The power of experienced iNaturalist users in enhancing ant ID accuracy

User experience, measured as the total number of IDs made across the dataset, proved to be a good predictor of the precision and accuracy of observations on iNaturalist. It was calculated as the average number of IDs from all contributing identifiers for each observation (*average_experience*). The Random Forest revealed that contributions from users with a higher number of identifications had a relevant impact on the consensus identifications, improving both precision and accuracy across all taxonomic categories studied and explaining a considerable amount of the variability in the models. Specifically, it was the second most influential variable in predicting precision and the first in accuracy.

To explore whether users with higher numbers of IDs individually produced more accurate identifications compared to less active or novice identifiers, users were divided into three activity levels based on their number of identifications: low activity (1–5 IDs), medium activity (6–20 IDs), and high activity (21+ IDs). Users were not evenly distributed across activity levels. Most users (142) fell into the low activity group, while fewer were in the medium (23) and high activity (13) groups. For this analysis, only observations with species-level identifications were selected (216 observations), and each user’s identification was compared against the expert’s. Results showed that the accuracy of species-level identifications varied across different user activity levels (Graph 6). The high activity group achieved the highest average accuracy (60.21%, SD = 24.67%), followed by the medium activity group (42.73%, SD = 20.30%) and the low activity group (34.88%, SD = 42.90%). A linear regression test revealed that the relationship was significant (p-value of 0.0277).

In conclusion, experienced users produced more accurate species-level identifications, and their participation in observations significantly improved precision and accuracy across all taxonomic levels in the final community consensus.

**Graph 6.**
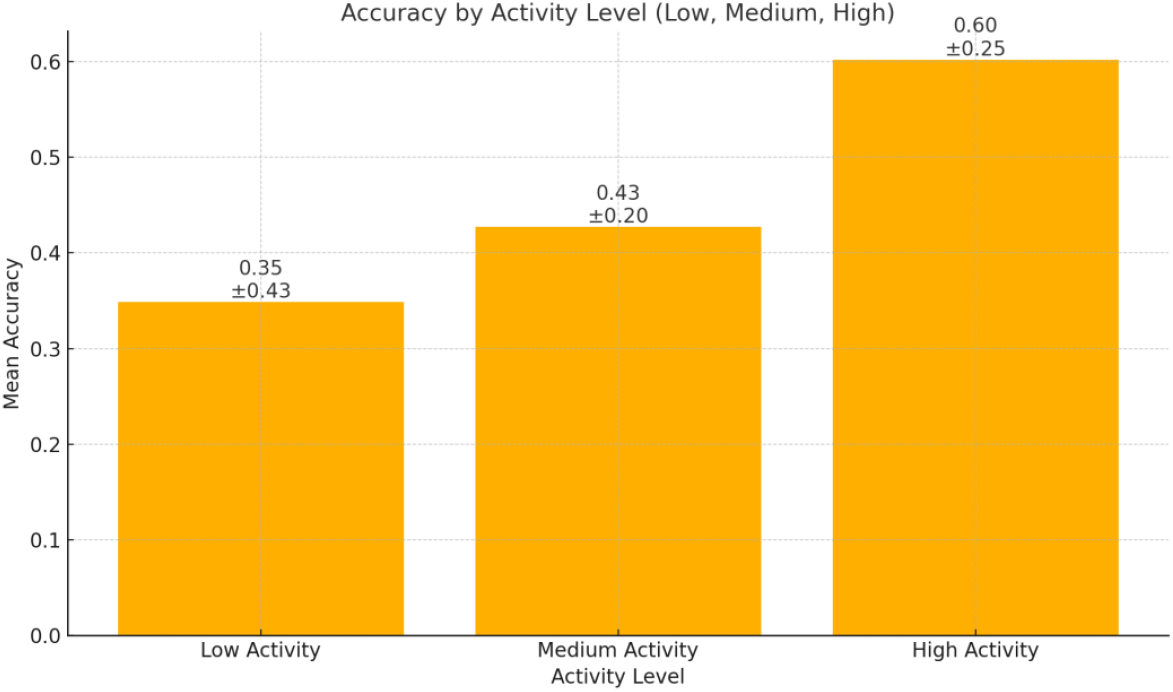
Accuracy of ID’s produced by identifiers of different experience levels (<10 IDs, 10-20 IDs, >20 IDs)

### 5. Familiar faces: how common ant species boost (and skew) ID accuracy

The study identified the total number of observations for each taxonomic unit (*total_observations*) as one of the most significant variables, explaining a great percentage of variability in accuracy and precision in the iNaturalist dataset. Taxa with more observations ranked with high accuracy compared to less common ants in the dataset. A closer look at the most observed ants revealed some common factors: they are conspicuous taxa, easy to spot in the field and common in the region of interest. For instance, the most common species, *M. bouvieri*, is conspicuous and large-sized, and the second one, *C. scutellaris*, is highly distinctive due to its bicolored red and black body. Precision in these species was 69.12% and 100.00%, respectively. Together, these two species accounted for 67.13% of the dataset (145 observations), significantly enhancing overall precision but also introducing a bias favoring easily recognizable taxa.

To further explore this bias, species were grouped based on their observation frequency, and individual and overall precision were calculated for each group (<10 observations, 10–20 observations, >20 observations). Commonly recorded species exhibited an accuracy of 82.07%, while moderately and infrequently recorded species achieved 61.29% and 32.50%, respectively (Table 6). A chi-squared test confirmed these differences were significant (χ^2^ = 37.90, p < 0.0001), suggesting a decline in community accuracy as observation frequency decreases. *M. ibericus* stood out as an outlier within its group, displaying a notably low precision (7.69%) despite being moderately observed. This is partly due to a recent nomenclatural change: *M. ibericus* is now the valid name for what was previously identified as *Messor structor* in Iberia (Steiner et al., 2018). With this taxonomic update, precision for *M. ibericus* improved to 23.07%.

**Table 6.** Species grouped by their total number of observations in the dataset, with both group average and individual precision. *Considering the recent nomenclatural change.

| Group | Species | Group precision | Precision by species |
| --- | --- | --- | --- |
| More than 20 observations | *Messor bouvieri*, *Crematogaster scutellaris*, *Pheidole pallidula* | 81.77% | *Messor bouvieri* (n=68, 69.0%), *Crematogaster scutellaris* (n=56, 100.0%), *Pheidole pallidula* (n=21, 76.0%) |
| 10-20 observations | *Linepithema humile*, *Messor ibericus* (13) | 53.85% | *Linepithema humile* (n=18, 100.0%), *Messor ibericus* (n=13, 23.07%*) |
| <10 observations | *Camponotus ruber*, *Camponotus barbaricus*, *Cardiocondyla mauritanica*, *Colobopsis truncata*, *Crematogaster laestrygon*, *Aphaenogaster iberica*, *Camponotus lateralis*, *Lasius balearicus*, *Messor capitatus*, *Myrmecina graminicola*, *Nylanderia vividula*, *Tetramorium bicarinatum*, *Tetramorium semilaeve* | 40.39% | *Camponotus ruber* (n=5, 100.0%), *Camponotus barbaricus* (n=4, 75.0%), *Cardiocondyla mauritanica* (n=2, 0.0%), *Colobopsis truncata* (n=2, 50.0%), *Crematogaster laestrygon* (n=2, 100.0%), *Aphaenogaster iberica* (n=1, 0.0%), *Camponotus lateralis* (n=1, 100.0%), *Lasius balearicus* (n=1, 0.0%), *Messor capitatus* (n=1, 0.0%), *Myrmecina graminicola* (n=1, 0.0%), *Nylanderia vividula* (n=1, 0.0%), *Tetramorium bicarinatum* (n=1, 100.0%), *Tetramorium semilaeve* (n=1, 0.0%) |

Infrequently recorded species exhibited a wide range of precision values, from 100% for *C. ruber* (5 observations) to 0% for *Cardiocondyla mauritanica* (2 observations). At the genus level, a similar pattern emerged (Graph 7). Commonly recorded genera, such as *Crematogaster* (98.53% accuracy, 68 observations), *Camponotus* (95.65%, 23 observations), and *Messor* (94.85%, 97 observations), displayed high accuracy rates. Moderately recorded genera, including *Linepithema* (100%, 18 observations) and *Tetramorium* (83.33%, 12 observations), also performed relatively well. However, uncommon genera like *Cardiocondyla* (2 observations) and *Myrmecina* (1 observation) demonstrated poor accuracy (both 0%), pointing to the difficulty of identifying some rarely encountered species in the Balearics. This overrepresentation of commonly observed taxa created an “easy win” effect, which inflated the perceived identification capacity for rarer species within the dataset.

To assess whether the behavior of identifiers differed between familiar and rare species, I compared the average number of identifications per observation for species with more than 20 observations, representing 48.33% of the dataset, to those with fewer observations. Notably, the most frequently recorded species received an average of 4.63 identifications per observation, slightly higher than the

4.39 average for less common ants. A t-test showed no statistically significant difference (p = 0.193), suggesting that the community’s level of engagement remained relatively consistent across species, regardless of their observation frequency.

**Graph 7.**
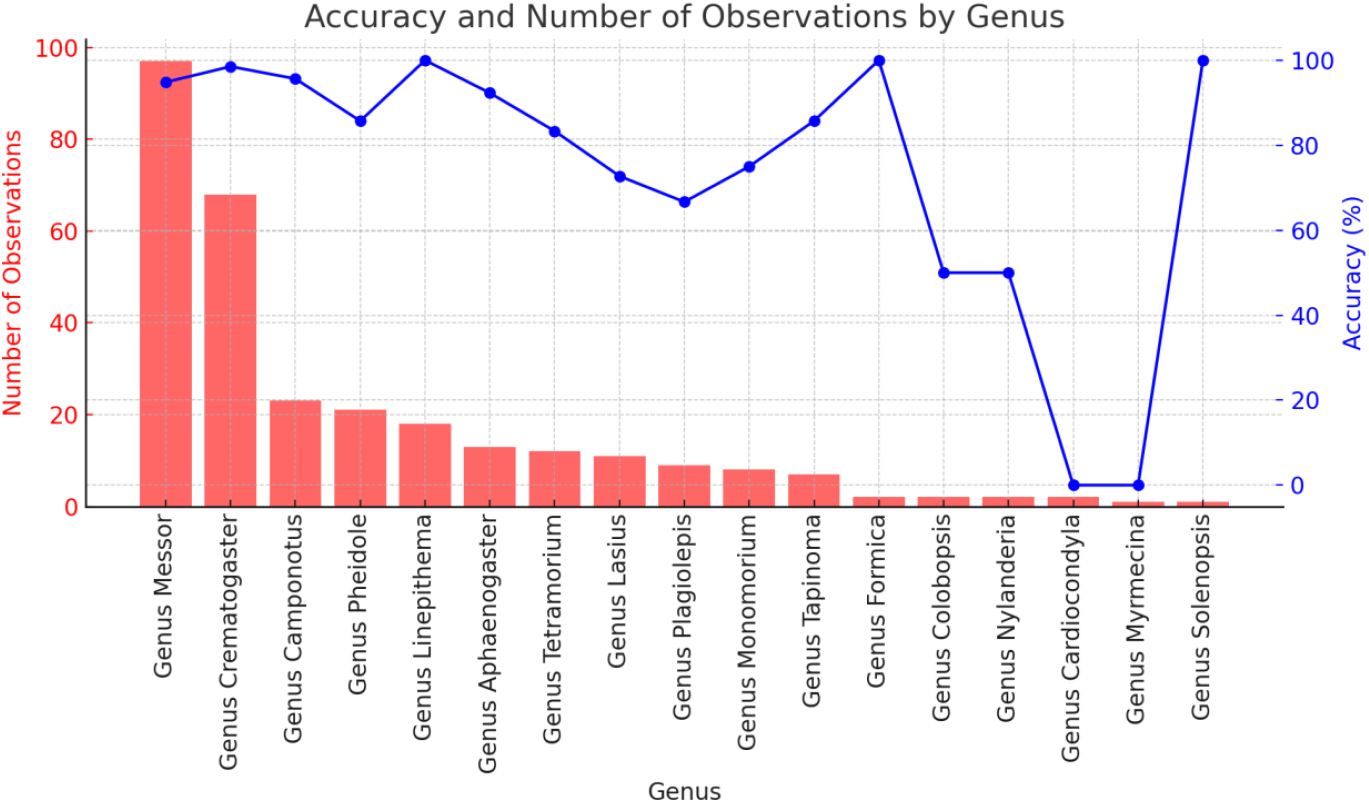
The graph illustrates the relationship between the number of observations and the accuracy of genus-level IDs.

### 6. ID 1 category: how early-stage classification impacts consensus

The taxonomic level at which the first identification (*ID1_category*) was proposed played a key role in the accuracy and precision of observations on iNaturalist, suggesting that early-stage classification had a significant impact on the likelihood of reaching a correct consensus. The analysis revealed that when the ID was made at finer taxonomic levels (species or genus), the final precision and accuracy of the observation were significantly higher than when ID1 was placed at broader levels (subfamily or family) (Graph 8, Graph 9). Specifically, when the first ID was at the species level, the final precision and average accuracy reached 77% and 92.7%, respectively, whereas when it was at the genus level, precision and average accuracy dropped to 54.9% and 72.7%. For subfamily-level initial IDs, accuracy fell below 50%, suggesting that overly broad initial classifications decreased the probability of a correct refinement by the community. The exception was at initial IDs at the family level, where precision and average accuracy showed better results than in previous categories, with 63.2% and 76.4%, respectively.

**Graph 8.**
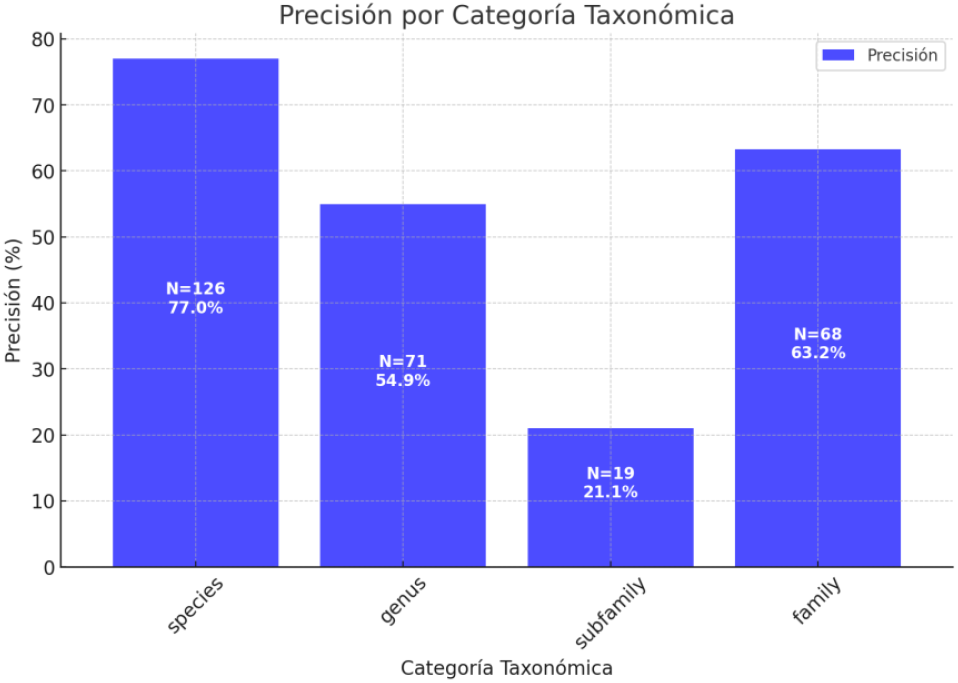
Final precision of observations grouped by the taxonomic level of the first identification.

**Graph 9.**
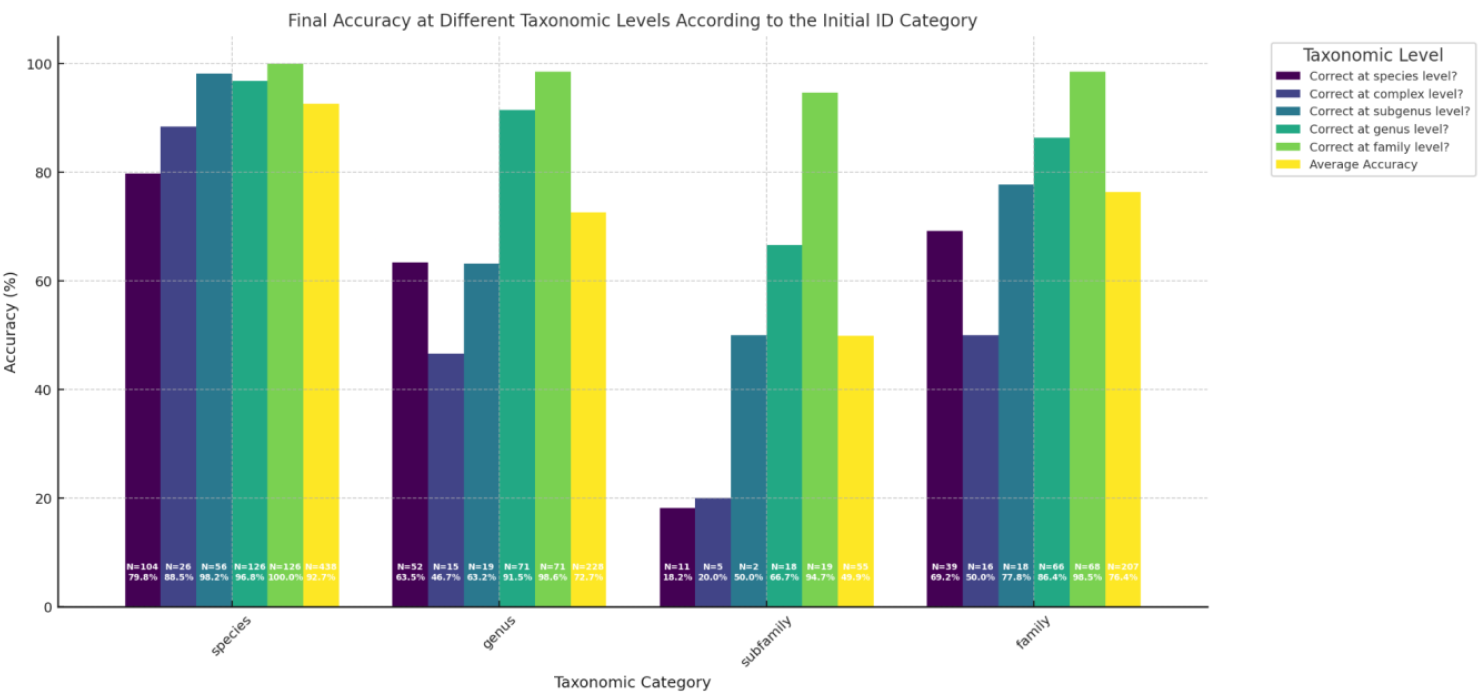
Final accuracy of observations (across multiple taxonomic categories) grouped by the taxonomic category of the first identification.

### 7. Is more always better? Changes in accuracy with increasing ID effort

Community effort was evaluated using three main variables: the total number of identifications (*total_IDs*), the number of identification agreements (*identification_agreements*), and the number of unique identifiers (*unique_identifiers_count*). Given that a higher number of identifiers generally leads to more IDs and agreements, these variables were expected to be highly correlated (see Section 1). A fourth variable, the number of identification disagreements (*identification_disagreements*), showed no significant correlation with the others and contributed minimally to the variability in precision and accuracy, likely because they were resolved through community consensus.

To examine the relationship between ID effort and accuracy, observations were grouped based on their number of IDs (ranging from 2 to 9). One observation with 10 IDs was excluded because of its outlier effect, and no observations had only one ID. The results revealed a clear positive trend: accuracy increased as the number of IDs grew (Graph 10). At lower ID counts (e.g., 2–3 IDs), accuracy ranged from 60% to 90% for both species and genus levels. When ID counts reached 6–9, accuracy improved substantially, with genus-level accuracy increasing more gradually than species-level accuracy. Precision followed a similar trend, starting at approximately 60% for observations with low ID counts and reaching 100% as the number of IDs increased (Graph 11).

Overall, this analysis emphasizes the relevance of collective participation in iNaturalist for enhancing both taxonomic accuracy and precision. The findings demonstrate that a higher number of identifications significantly improves accuracy at the species and genus levels, reinforcing the value of multiple contributions in achieving more reliable taxonomic classifications.

**Graph 10.**
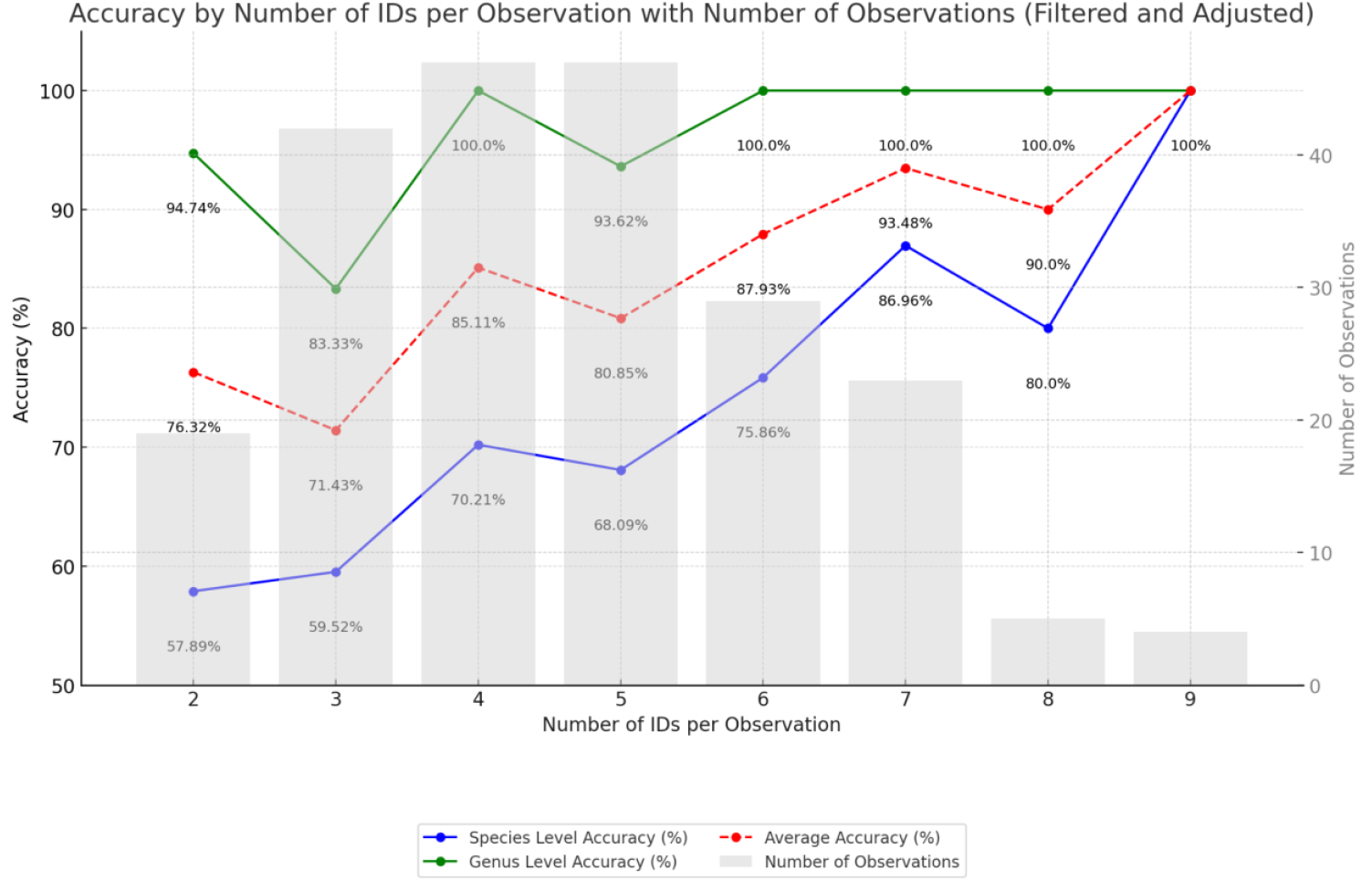
Changes in accuracy by number of identifications per observation.

**Graph 11.**
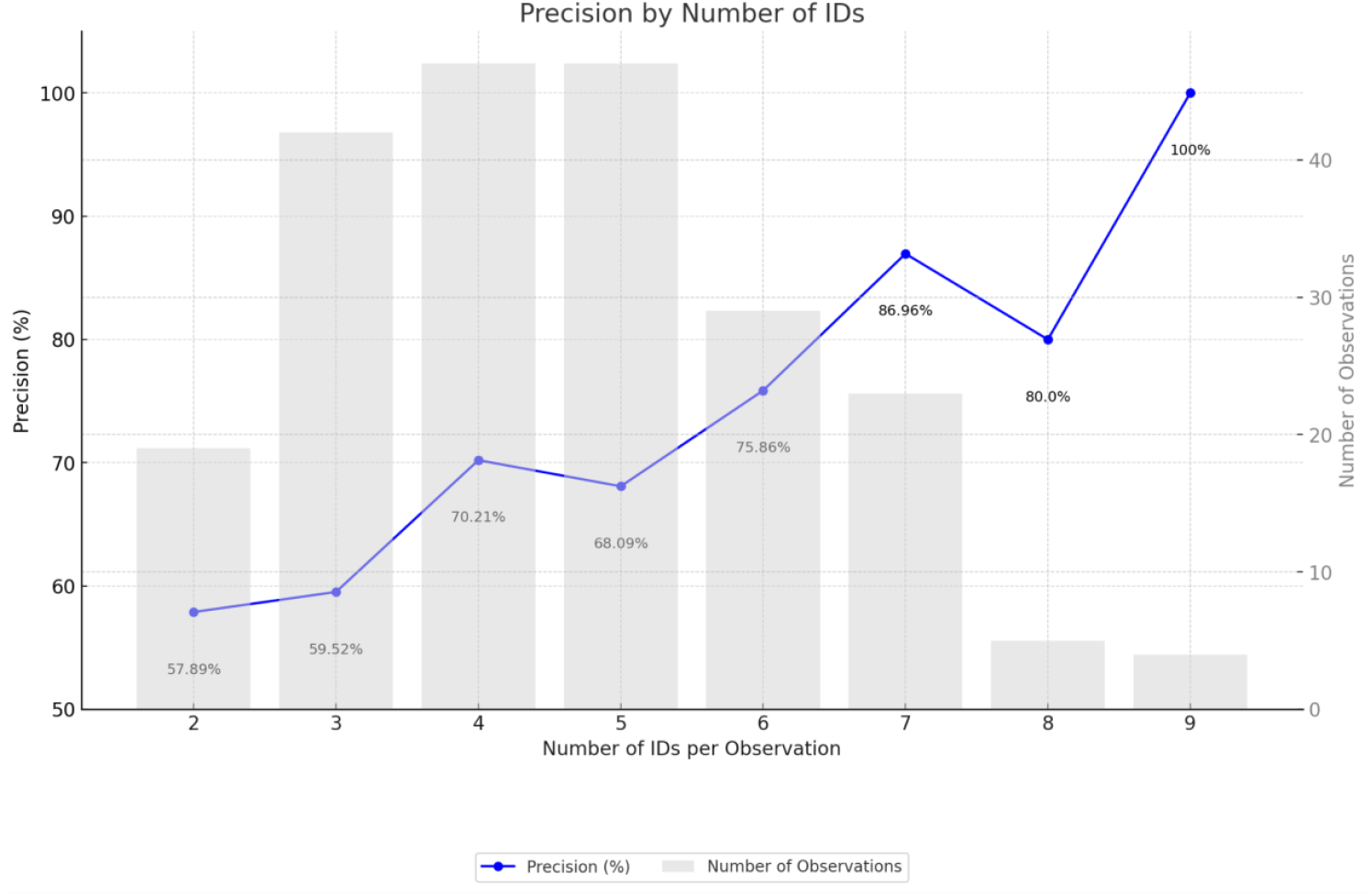
Changes in precision by number of identifications per observation.

### 8. Time Will Tell: Temporal Trends in Identification Accuracy from 2018 to 2024

The passage of time demonstrated a small degree of precision and accuracy variance in the analysis. The study examined trends in taxonomic accuracy at different hierarchical levels—species, genus, sub-genus, and subfamily—from 2018 to 2024. For each of the 1371 unique IDs in the 300 observations, I classified them by the year they were made and compared their accuracy at different taxonomic levels whenever they were available. The average accuracy of IDs for each year was calculated and is presented in Graph 12. The analysis revealed a clear upward trend in the accuracy of ant identifications on iNaturalist from 2018 to 2024 across all taxonomic levels. Genus-level accuracy consistently remained high, increasing from 78.3% in 2018 to 96.5% in 2024. Similarly, subfamily-level accuracy improved from 83.3% to 97.6% over the same period. Species-level accuracy, although initially lower at 65.0%, exhibited the most substantial improvement, reaching 80.2% by 2024. The overall average accuracy across all taxonomic levels also increased from 68.5% in 2018 to 86.4% in 2024. These results indicate a notable improvement in ant identification accuracy on iNaturalist from 2018 to 2024, highlighting the platform’s growing data reliability.

**Graphic 12.**
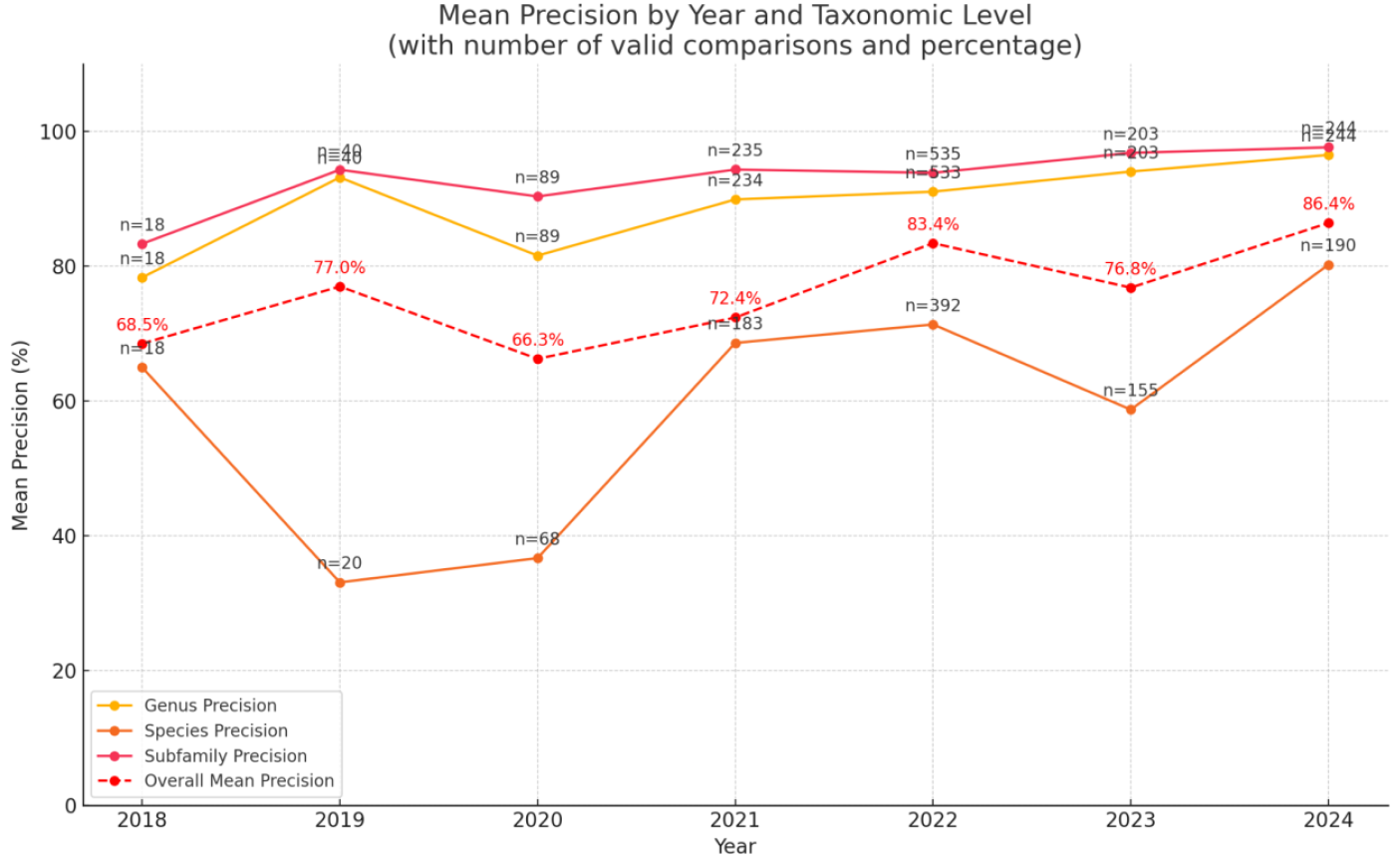
Yearly average accuracy of ant identifications by taxonomic level. The red dashed line represents the average accuracy across all considered taxonomic levels for each year. The number of valid comparisons (n) used to calculate the accuracy for each year is indicated above the corresponding points.

### 9. Artificial Intelligence at work: strengths and limitations

Beginning in 2017, the iNaturalist Computer Vision system is integrated in the platform as an AI-based taxonomy identifier, trained by user-uploaded observation images. Over time, continuous updates, expanded taxonomic coverage, and user feedback have gradually enhanced the system’s accuracy. Despite these improvements, the system’s precision across different taxonomic levels—species, genus, and subfamily—and its concordance with expert identifications remain important areas for further investigation.

Several key operational aspects of the AI system in iNaturalist are worth noting. First, when uploading their photos, users are shown an AI-generated taxonomic suggestion, which they can then accept or reject. Only taxa included in the AI training model can receive AI-based suggestions, while untrained taxa do not receive automated suggestions. AI suggestions can span from species-level identifications to family-level identification. For example, in our dataset, the system suggested “Family Formicidae” for 11.67% of observations (35/300), followed by “*Crematogaster scutellaris*” (11.33%, 34/300) and “Genus *Messor*” (8.33%, 25/300). Finally, identifiers can offer new identifications to an observation, potentially informed by the AI suggestion. However, the platform does not track whether identifiers adhere to the AI’s suggestion, which highlights a gap in understanding the AI’s influence on user behavior.

In our analysis, we focused on two variables related to AI use in ant identification on iNaturalist: *AI_suggestion*, which indicates whether an AI-generated suggestion was provided initially as the first ID, and *AI_possibility*, which reflects the inclusion of taxa in the AI’s training model. Their overall impact on identification precision and accuracy was limited in the Random Forest models. Within the dataset of 300 observations, 196 (65.33%) included an AI-generated suggestion, whereas 104 (34.67%) did not. When observers adopted the AI-generated suggestion, the taxonomic resolution aligned with the expert’s identification in 58 of 196 cases (29.59% precision). Among observations with AI suggestions, taxa were correctly identified at the species level in 54.1% of cases, identified correctly at a higher taxonomic level in 26.5%, and identified incorrectly at the species level in 19.4%. When focusing on taxa explicitly included in the iNaturalist Computer Vision model, noticeable gains in precision were observed. For taxa outside the AI’s training scope (Possibility of AI = “No”), only 25 out of 65 observations (38.46%) were precise, whereas 40 observations (61.54%) were imprecise. Conversely, for taxa within the AI’s training scope (Possibility of AI = “Yes”), precision rose to 167 out of 235 observations (71.06%), with 68 observations (28.94%) imprecise. This analysis points to the necessity of expanding the AI training dataset to improve identification accuracy, particularly at finer taxonomic resolutions.

### 10. Out of focus? The limited role of photo quality in ant ID precision

The impact of photo quality on the ant identification models was found to be minimal. Ant species identification often relies on subtle morphological characters, such as the shape of the peduncle or the pilosity density of different parts of the body, which are often indistinguishable in photographs regardless of their quality. However, for commonly recorded species like *M. bouvieri* or *C. scutellaris*, medium or low-quality photos are often sufficient for a confident identification, as their general appearance is distinct from other similar taxa in the Balearics. The already discussed familiarity of the user community with these common species appears to reduce the necessity for high-quality images. When a species cannot be confidently identified, users tend to assign a higher taxonomic rank (e.g., genus or family). This may explain why photo quality has a greater influence at the genus level, where less intricate details are sufficient for identification. Additionally, no statistically significant difference was found between the number of observations across taxonomic units and the photo quality (p = 0.552), suggesting that photo quality is not biased toward more or less common species.

### 11. Newcomers and old Threats: insights on the exotic ants in the Balearic Islands

Several exotic ant species have been documented as established in the Balearic Islands (Gómez & Espadaler, 2006; Díaz-Calafat & Fortis, 2022; Arcos & Alarcón, 2024). In the analyzed dataset, exotic ants accounted for 30 observations across seven species, representing 10% of the total dataset. The species *L. humile*, commonly known as the Argentine ant, constituted the majority of these observations (60% of the total exotic ant observations). Recognized globally as one of the most invasive species, L. humile has caused significant ecological disruptions, including reductions in biodiversity and alterations to community structures (Angulo et al., 2024). It was introduced to Mallorca in the 1950s, and has since spread across the three major islands, with devastating effects on native ant populations (Gómez & Espadaler, 2005).

In addition to *L. humile*, other exotic ant species, such as *C. mauritanica, N. vividula*, and *A. iberica*, have been established in the Balearic Islands for decades. More recent introductions, including *C. barbaricus* and *T. bicarinatum*, have also been reported (Díaz-Calafat & Fortis, 2023; Arcos & Alarcón, 2024). Notably, one of the earliest records of *T. bicarinatum* originated from the iNaturalist platform. Meanwhile, *C. barbaricus* has been recorded in Mallorca at two localities, and at least five new iNaturalist observations suggest its successful local spread in one of them (Andratx). A significant discovery involves the *F. rufibarbis* complex, with observations of 2022 and 2024 reporting its presence in Mallorca, confirming the first records of this species in the islands. These observations, geolocated near Parc de sa Riera in Palma, Mallorca (https://www.inaturalist.org/observations/118939446; https://www.inaturalist.org/observations/226375396), indicate its presence in an anthropogenic environment. The absence of historical records for this species in the region (Gómez & Espadaler, 2006) strongly suggests it is an exotic species. Differentiation between the two possible candidates of this cryptic species group, *Formica rufibarbis* and *Formica clara*, requires detailed morphometric analysis, as they are visually indistinguishable (Seifert, 2007). The spatial proximity of these observations raises uncertainty about whether they represent a single colony or multiple colonies.

Between 7.6% and 17.4% of annual observations involved exotic ant species, with a peak in 2021. Despite the discovery of three significant new exotic species in recent years, the data do not indicate a consistent upward trend in their overall presence. On the contrary, the proportion of observations involving exotic and invasive ants, including the dominant *L. humile*, has decreased over time. Nevertheless, the establishment and spread of exotic species in the Balearics remain an ongoing threat to native biodiversity. While *L. humile* remains the most prevalent exotic species, recent introductions and the spread of *T. bicarinatum, C. barbaricus*, and the *F. rufibarbis* complex highlight the need for continued vigilance.

**Graph 13.**
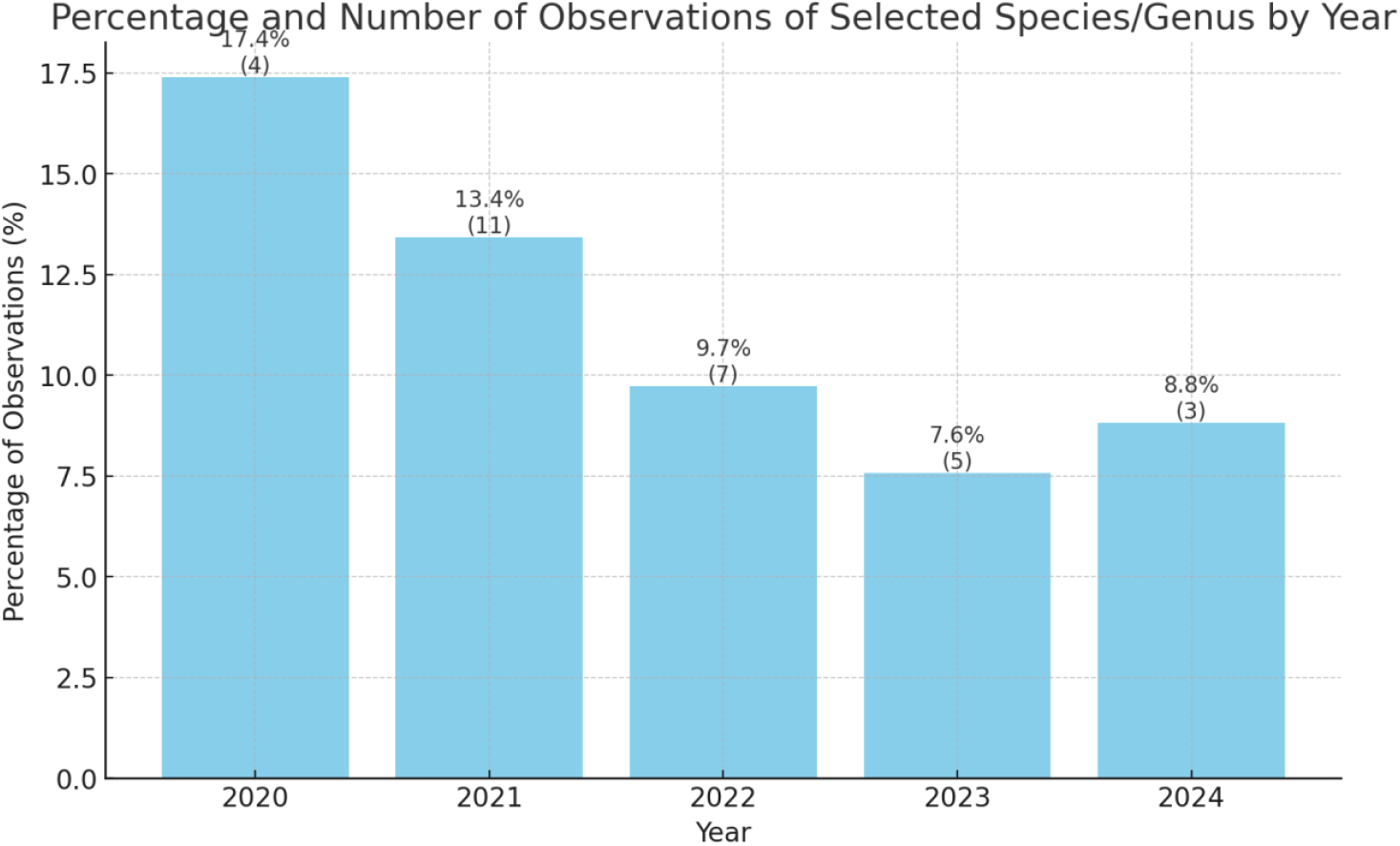
Changes in percentage of exotic ant observations by the total observations uploaded each year.

### 12. Additional results

There are other interesting results from the analysis of iNaturalist observations of ants from the Balearic Islands. For the first time, the undescribed queen of *Lasius balearicus* Talavera et al., 2015 —an endemic and critically endangered species— was photographed (see observation: https://www.inaturalist.org/observations/193781076).

Images of workers from the *T. caespitum* complex account for 30% of the observations. These are likely *Tetramorium immigrans*, an exotic ant species not yet reported on the islands. However, morphometric analysis is required to confirm this ID and differentiate it from other members of the complex (Steiner et al., 2018).

Although not analyzed in this paper, some observations of ants on iNaturalist correspond to reproductive individuals (males or queens). However, their representation seems to be relatively scarce, and its identification precisions probably significantly lower than those of workers.

## Discussion

This study demonstrates both the strengths and limitations of iNaturalist for ant identification in the Balearic Islands. While overall precision of community identifications was moderate (64%), accuracy improved significantly at broader taxonomic levels, reaching 90.91% for genus and 96.31% for subfamily. Species-level accuracy (69.91%) was comparable to previous studies on diverse taxa, such as terrestrial molluscs (54% accuracy) (Barbato et al., 2021), and vascular plants on iNaturalist, where species-level precision reached 72.3% and improved to 85.6% at the genus level (Wenk et al., 2024). In contrast, a study on lichens reported a 70% misidentification rate, highlighting the challenges of species-level identification in certain groups (Munzi et al., 2023).

Despite the well-documented ant fauna of the Balearic Islands, where 67 species are known (Arcos & García, 2024), only 26 taxonomic units were recorded in this study, with 18 species identified at the species level. The dataset was strongly biased toward a few common species, with the top 10 species accounting for 95.67% of all observations and nearly 50% of observations belonging to just two species, *M. bouvieri* and *C. scutellaris*. This underrepresentation of taxonomic diversity aligns with previous studies that show citizen science platforms often overrepresent conspicuous and easily identifiable taxa (Di Cecco et al., 2021; Wenk et al., 2024). The presence of species that could not be identified to species level due to photographic limitations further highlights the challenge of documenting the full diversity of ants in the region.

User experience played a very significant role in data quality of ant observations. High-activity users demonstrated greater expertise in individual identifications, and their participation also improved overall community accuracy. This aligns with findings that frequent contributors provide higher-quality data (Kosmala et al., 2016; Campbell et al., 2023). However, unlike other studies where observers greatly outnumber identifiers (Campbell et al., 2023), this study found a proportion of 1:1.71 observers to identifiers, suggesting a more balanced distribution of effort among users. This balance may contribute to the relatively high identification accuracy observed, as a greater proportion of observations received multiple independent evaluations.

Additionally, the taxonomic resolution of the initial identification was found to be an important factor explaining accuracy and precision, as observations where the first identification was placed at a finer taxonomic level (e.g., species) were more likely to reach a correct final consensus than those starting at broader levels. For instance, when the first identification was made at the species-level, the final precision reached 77%, while for observations where the first identification was at the genus-level, precision dropped to 54.9%.

The role of community effort was also significant, as the accuracy of identifications increased with the number of submitted IDs. Observations with only 2–3 IDs had an accuracy ranging from 60% to 90%, whereas those with 6–9 IDs showed a marked improvement, particularly at the species level. Precision followed a similar trend, starting at approximately 60% for observations with low ID counts and reaching 100% as the number of IDs increased. Observations with a higher number of identifiers tended to have a greater likelihood of reaching a correct consensus, reinforcing the importance of collective participation in citizen science.

Temporal trends in identification accuracy showed a steady improvement from 2018 to 2024. Genus-level accuracy consistently remained high, increasing from 78.3% in 2018 to 96.5% in 2024, while sub-family-level accuracy rose from 83.3% to 97.6%. The most notable increase was observed at the species level, where accuracy improved from 65.0% in 2018 to 80.2% in 2024. The overall average accuracy across all taxonomic levels also increased from 68.5% in 2018 to 86.4% in 2024, indicating a progressive improvement in ant identification reliability on iNaturalist

AI-generated suggestions were present in 65.33% of observations. When taxa were included in the AI training dataset, precision was 71.06%, but for taxa outside the model, it dropped to 38.46%, demonstrating AI’s dependence on the availability of training data. While AI improved accuracy for frequently observed species, it struggled with rarer or underrepresented taxa, similar to previous studies where AI models underperformed for species not well-represented in training datasets (Campbell et al., 2023). Expanding AI datasets to include a broader range of species could enhance its reliability, particularly for cryptic or less frequently recorded taxa.

Despite expectations, photographic quality had a limited impact on accuracy. Many species were correctly identified even from low-quality images, suggesting that for certain taxa, general morphology is sufficient for identification. However, this does not hold for cryptic species, which require microscopic or chemical analysis for definitive identification. Prior research has already shown that accuracy for cryptic or difficult-to-identify taxonomic groups tends to be low, even when species are theoretically identifiable from photographs alone (McMullin & Allen, 2022). This suggests that while iNaturalist can provide valuable data for some taxa, its effectiveness is constrained by taxonomic and methodological limitations.

Citizen science proved valuable for detecting exotic species, with 10% of observations representing introduced ants, including *L. humile* and the first records of the *F. rufibarbis* complex in Mallorca. This supports the role of citizen science in tracking invasive species, as observed in other studies (Encarnação et al., 2021). These results indicate that citizen science platforms can be effective for biodiversity monitoring but also highlight the need for targeted efforts to document underrepresented taxa.

This study also provides valuable insights at multiple taxonomic levels, rather than focusing solely on species-level accuracy, as other previous studies have done. Analyzing accuracy at the genus and sub-family levels offers additional taxonomic information that can improve species distribution models and biodiversity assessments.

## Conclusions

This study shows that iNaturalist can provide reliable data for ant identification, as illustrated by the Balearic Islands case study, but its reliability depends on multiple factors. While overall precision was moderate, accuracy increased at broader taxonomic levels, and user experience, community participation, and taxon familiarity played key roles in determining the correctness of identifications. The results highlight that citizen science contributions are most reliable for common and easily recognizable species, which make up most of the dataset, while rare or morphologically complex taxa remain challenging to identify accurately. Artificial intelligence algorithms enhanced identification accuracy when species were already included in the training dataset, but their effectiveness diminished for less represented taxa. Despite its limitations, iNaturalist proved to be a useful tool for biodiversity monitoring, including the detection of exotic ant species. Continued improvements in AI models, increased expert participation, and enhanced community engagement could further strengthen the reliability of citizen-generated data. Future studies should focus on refining methods to improve accuracy, particularly for cryptic or underrepresented species, ensuring that citizen science remains a robust resource for ecological research and conservation.

